# Perturbations of the T-cell immune repertoire in kidney transplant rejection

**DOI:** 10.1101/2022.08.24.505187

**Authors:** Tara K. Sigdel, Paul A. Fields, Juliane Liberto, Izabella Damm, Maggie Kerwin, Jill Hood, Parhom Towfighi, Marina Sirota, Harlan S. Robins, Minnie M. Sarwal, the CMV Systems Immunobiology Group

**Author notes:** **Consortia Authorship**, CMV Systems Immunobiology Group. **Corresponding Author**, Minnie M. Sarwal, MD, PhD, Professor in Residence, Surgery, Medicine, Pediatrics, UCSF, Medical Director, Pancreas-Kidney Transplant, UCSF, 513 Parnassus Ave, Med Sciences Bldg., Room S1268, San Francisco, CA 94143.

## Abstract

In this cross-sectional and longitudinal analysis of mapping the T-cell repertoire in kidney transplant recipients, we have investigated and validated T-cell clonality, immune repertoire chronology at rejection, and contemporaneous allograft biopsy quantitative tissue injury, to better understand the pathobiology of acute T cell and antibody-mediated kidney transplant rejection. To follow the dynamic evolution of T-cell repertoire changes before and after engraftment and during biopsy-confirmed acute rejection, we sequenced 323 peripheral blood samples from 200 unique kidney transplant recipients, with (n=100) and without (n=100) biopsy-confirmed acute rejection. The results of these studies highlight, for the first time, that patients who develop acute allograft rejection, have lower (p=0.01) T cell fraction even before transplantation, followed by its rise after transplantation and at the time of acute rejection accompanied by high TCR repertoire turnover (p=0.004). Acute rejection episodes occurring after the first 6 months post-transplantation, and those with a component of antibody-mediated rejection, had the highest turnover; p=0.0016) of their TCRs. In conclusion, further prospective validation studies are needed to evaluate the clinical utility of peripheral blood TCR analysis for both pre- and post-transplant immune risk assessment and prediction of different mechanisms of graft rejection.

## Introduction

Transplanted kidneys fail due to both immune and nonimmune causes including primarily anti-donor alloimmunity via recruitment of activated T cells, but also due to other contributing factors such as activation of innate immunity by triggers such as cold ischemia time, donor co-morbidities, heterologous immunity from infections, and drug toxicities [1]. Recent developments in T-cell receptor (TCR) sequencing allow for analysis of alloreactive human T-cell populations both before and after engraftment [2–4]. Accurate and high-throughput sequencing of the third complementarity-determining region (CDR3) of the TCRβ chain, which is the crucial segment for antigen specificity, allowed us to profile the T-cell repertoire in peripheral blood sample, coincident with the occurrence of acute renal transplant rejection on a paired allograft biopsy, and to evaluate if rejection could be predicted prior to histological evaluation by peripheral blood sampling before transplantation and if specific changes in the TCR repertoire can define different phenotypes of acute rejection, such as antibody mediated rejection[4].

To assess the evolution of the T-cell immune repertoire in peripheral blood samples and their patterns in the context of allograft phenotypes, we compared peripheral blood samples from demographically matched renal transplant recipients who had either protocol or clinically indicated biopsies and paired blood samples for immune repertoire analysis; group comparisons were conducted between patients with histologically clean protocol biopsies (designated as the stable group (STA)) and histologically confirmed acute rejection as graded by the Banff classification on either protocol or indicated biopsies (designated as the acute rejection group (AR) [5, 6]. Recent exploratory studies tracking peripheral blood T cells using TCR sequencing after kidney transplantation [5, 7–10] have observed low-frequency immune repertoires alterations with small sample sizes [11, 12]. In children and young infants, a diverse B-cell pre-transplant repertoire, correlated with post-transplant rejection risk, but no differences were found in the TCR immune repertoire. [13]. Having found no significant changes in the TCR repertoire in acute rejection in pediatric recipients of renal allografts, in this study, we evaluated the TCR immune repertoires in a large cohort of adult renal transplant patients so as to be able to discern low frequency changes between T-cell clones and transplant time, histologically confirmed acute rejection episodes, and any specific variations with T cell or antibody mediated rejections.

## Materials and methods

### Sample collection

The objective was to assess the evolution of the T-cell immune repertoire in peripheral blood samples and their patterns in the context of allograft phenotypes. For this, the study included biobanked biopsy-matched total blood samples (n=323) collected by an IRB approved research protocol at the University of California San Francisco (Study ID-#14-13573) and selected from a biobank of over 3000 blood samples collected from kidney transplant recipients. The blood samples selected for this study included pre-transplant baseline samples from 100 renal allograft recipients who had biopsy-confirmed AR, inclusive of both T cell-mediated rejection (TCMR) and antibody-mediated rejection (ABMR); 100 renal allograft recipients had STA biopsies and graft function. All blood samples were submitted for sequencing using the ImmunoSEQ Assay (Adaptive Biotechnologies, Seattle, WA). For STA patients, 50 subjects had a 6-month post-transplant sample at the time of management (protocol) biopsy and none of them went on to develop AR over the next year of follow-up. For AR patients, 50 subjects had a blood sample at the time of AR, and 39 of those patients had an additional 6-month management biopsy blood draw. All subjects were enrolled following IRB approval and had informed consent. Study subjects’ baseline characteristics are summarized in **Table 1**.

**Table 1.**
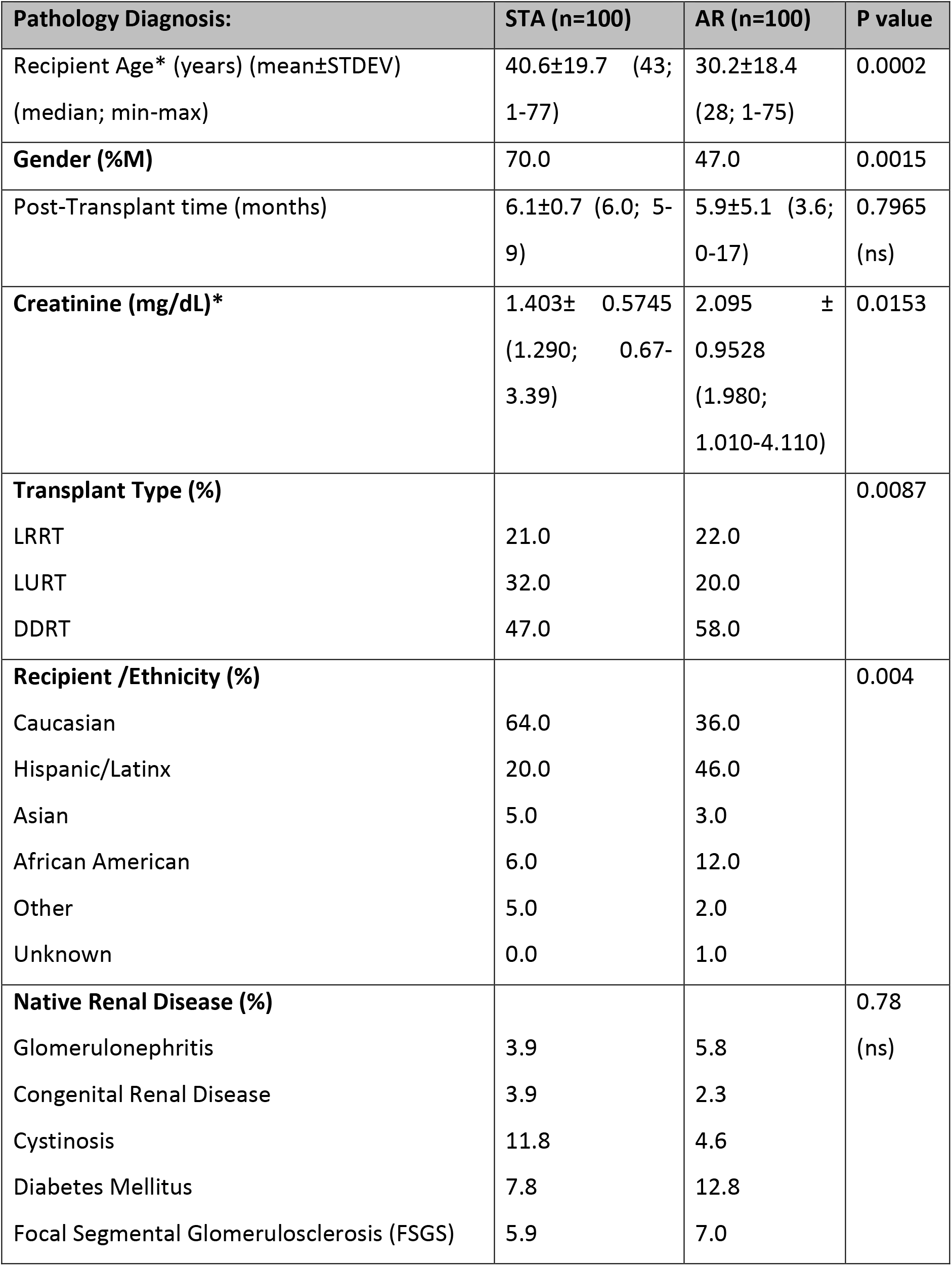
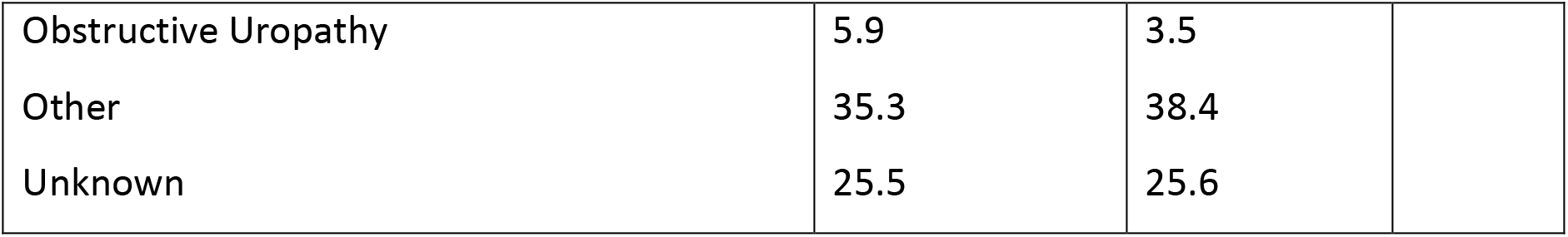
Demographic and Baseline Characteristics.

| **Pathology Diagnosis:** | **STA (n=100)** | **AR (n=100)** | **P value** |
| --- | --- | --- | --- |
| Recipient Age* (years) (mean±STDEV)<br>(median; min-max) | 40.6±19.7 (43;<br>1-77) | 30.2±18.4<br>(28; 1-75) | 0.0002 |
| **Gender (%M)** | 70.0 | 47.0 | 0.0015 |
| Post-Transplant time (months) | 6.1±0.7 (6.0; 5-9) | 5.9±5.1 (3.6;<br>0-17) | 0.7965<br>(ns) |
| **Creatinine (mg/dL)\*** | 1.403± 0.5745<br>(1.290; 0.67-3.39) | 2.095 ±<br>0.9528<br>(1.980;<br>1.010-4.110) | 0.0153 |
| **Transplant Type (%)** |  |  | 0.0087 |
| LRRT | 21.0 | 22.0 |  |
| LURT | 32.0 | 20.0 |  |
| DDRT | 47.0 | 58.0 |  |
| **Recipient /Ethnicity (%)** |  |  | 0.004 |
| Caucasian | 64.0 | 36.0 |  |
| Hispanic/Latinx | 20.0 | 46.0 |  |
| Asian | 5.0 | 3.0 |  |
| African American | 6.0 | 12.0 |  |
| Other | 5.0 | 2.0 |  |
| Unknown | 0.0 | 1.0 |  |
| **Native Renal Disease (%)** |  |  | 0.78<br>(ns) |
| Glomerulonephritis | 3.9 | 5.8 |  |
| Congenital Renal Disease | 3.9 | 2.3 |  |
| Cystinosis | 11.8 | 4.6 |  |
| Diabetes Mellitus | 7.8 | 12.8 |  |
| Focal Segmental Glomerulosclerosis (FSGS) | 5.9 | 7.0 |  |
| Obstructive Uropathy | 5.9 | 3.5 |  |
| Other | 35.3 | 38.4 |  |
| Unknown | 25.5 | 25.6 |  |

The matching allograft biopsy at the time of blood sampling was read by a single pathologist using semi-quantitative histological scores. Clinical AR was defined as an AR episode, associated with graft dysfunction, based on a greater than 20% rise in serum creatinine from baseline values, and confirmed through central pathological reading of the biopsies according to the updated Banff classification with semi-quantitative scoring for tubulitis (0, t1, t2), inflammation (0, I1, i2) and interstitial fibrosis and tubular atrophy (IFTA; 0, 1=<10% IFTA, 2= 10-50% IFTA, 3= >50% IFTA)[14]. All samples were collected at UCSF under IRB-approved protocols approved by The Human Research Protection Program (HRPP) of the University of California, San Francisco (UCSF) to allow analysis of bio-banked samples. CMV IgG status was measured on donor and recipient pairs and CMV PCR at 6 months post-transplant and with clinical suspicion in all the participants. All study subjects received antivirus prophylaxis until 6 months posttransplant. All patients or their guardians provided informed consent to participate in the research in full adherence to the Declaration of Helsinki. The clinical and research activities being reported are consistent with the Principles of the Declaration of Istanbul as outlined in the Declaration of Istanbul on Organ Trafficking and Transplant Tourism.

### Isolation of genomic DNA and T-cell receptor variable beta chain sequencing

Blood samples (4.5 ml) were collected into a 5 ml red top tube, gently inverted, and incubated at room temperature for 30 min until the clot was formed. The sample was then centrifuged at 2000 × *g* for 5 min, the upper layer of serum was then transferred to a cryotube, and the remaining clot was stored in the original tube at −80 °C until use. Genomic DNA from whole-blood clot was extracted using the QIAsymphony DSP DNA Midi Kit (Qiagen, Valencia, CA). The DNA was quantified with NanoDrop and the extracted DNA was stored in −80°C until use for sequencing of the CDR3 regions of human TCRβ locus using the immunoSEQ assay. Extracted genomic DNA was amplified in a bias-controlled multiplex PCR with a pool of primer pairs targeting the V and J genes, as well as primers targeting reference genes to quantitate the total nucleated cells in each sample. PCR products were sequenced on an Illumina NextSeq System. Sequences were collapsed and filtered in order to identify and quantitate the absolute abundance of each unique TCRβ CDR3 region for further analysis as previously described [2, 3, 15–18]. Twelve samples that had less than 500 total T cells were excluded from analysis due to insufficient DNA yields, resulting in 327 samples being included in the analyses (STA: 98 at baseline, 49 at 6M, AR: 92 at baseline, 43 at <6M, 35 at 6M, 49 at >6M AR time post-transplantation) (**Table 2**).

**Table 2:** Sequencing statistics. Mean (Min – Max)

| **Outcome / Time** | **Samples** | **Total T Cells** | **Unique Productive Rearrangements** | **T cell Fraction** | **Simpson Clonality** |
| --- | --- | --- | --- | --- | --- |
| STA\|D0 | 98 | 102208<br>(590-386883) | 54859<br>(539-231736) | 0.222<br>(0.001-0.557) | 0.044<br>(0.002-0.396) |
| STA\|6M | 49 | 68140<br>(2184-302113) | 29913<br>(1600-111518) | 0.161<br>(0.035-0.62) | 0.067<br>(0.009-0.516) |
| AR\|D0 | 92 | 69333<br>(525-470858) | 46094<br>(301-390454) | 0.167<br>(0.001-0.547) | 0.037<br>(0.003-0.268) |
| AR\|6M | 35 | 50139<br>(4137-142752) | 23577<br>(3099-127188) | 0.184<br>(0.02-0.341) | 0.047<br>(0.004-0.234) |
| AR\|AR<6mo | 25 | 29481<br>(3998-163414) | 19662<br>(2858-103045) | 0.069<br>(0.009-0.411) | 0.069<br>(0.01-0.309) |
| AR\|AR>6mo | 19 | 106176<br>(14229-218907) | 47528<br>(6418-144186) | 0.179<br>(0.024-0.459) | 0.059<br>(0.004-0.186) |
| AR\|AR | 5 | 45983<br>(29528-71191) | 22877<br>(21639-31523) | 0.134<br>(0.091-0.173) | 0.045<br>(0.017-0.104) |

*5 AR samples are excluded in longitudinal analysis as they failed QC*

TCRβ sequencing data that support the findings of this study have been deposited at the publicly available immuneACCESS platform at: https://clients.adaptivebiotech.com/pub/<DETERMINEDONACCEPTANCE>.

We compared TCRβ sequencing data from demographically matched renal transplant recipients who had either protocol or clinically indicated biopsies and paired blood samples for immune repertoire analysis; group comparisons were conducted between patients with histologically clean protocol biopsies (designated as the stable group (STA)) and histologically confirmed acute rejection as graded by the Banff classification on either protocol or indicated biopsies (designated as the acute rejection group (AR)

### Statistical analyses of TCRβ sequencing results

Simpson clonality was calculated on productive rearrangements using the following equation: 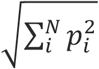 where *p_i_*, is the proportional abundance of rearrangement *i* and *N* is the total number of rearrangements [19]. Clonality values range from 0 to 1 and describe the shape of the frequency distribution: clonality values approaching 0 indicate a very even distribution of frequencies, whereas values approaching 1 indicate an increasingly asymmetric distribution in which a few clones are present at high frequencies.

TCFr, or the proportion of nucleated cells that are T cells, was computed using the number of T cells as determined by sequencing of productive TCRs and the number of nucleated cells by simultaneously amplifying a panel of reference genes present in all nucleated cells [19]. We determine both the total number of T cells and the total number of nucleated cells using our bias controlled PCR methodology[18]. Morisita Index was used to compare repertoires between longitudinal samples from the same individual. Morisita Index was calculated with the following formula: 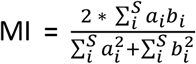. *a_i_*, equals the frequency of clone *i* in sample a, and *b_i_* equals the frequency of clone *i* in sample b. Morisita index takes into account sample overlap and change in clone frequencies with values that range from 1 (identical repertoires) to 0 (completely disparate repertoires) [20].

All statistical analysis was performed using standard packages in R version 3.6. Associations between groups were tested with either a Wilcoxon Rank Sum Test (two categorical groups) or a Kruskal-Wallis Test (three or more categorical groups). Pairwise comparisons after Kruskal-Wallis Tests were conducted with a Post-Hoc Dunn Test. Associations between two continuous variables were tested using Spearman’s Rank-Order Correlation. Associations between pairs of categorical groups were tested with a Fisher Exact Test.

## Results

The study cohort included a carefully selected set of peripheral blood samples obtained from demographically matched renal allograft patient groups with and without biopsy-confirmed acute renal allograft rejection[14], and with a contemporaneously obtained peripheral blood sample obtained prior to treatment intensification of graft rejection. These samples were selected from our larger biobank of over 3000 blood samples; we included 100 patients with that met definition of the STA group and 100 patients that met the definitions of the AR group, as described above. All patients had blood samples profiled prior to transplantation with the aim of evaluating changes if changes in the TCR immune repertoire could predict acute rejection after transplantation. In 50% of the patients, paired blood samples were also collected and profiled at the time of the post-transplant biopsy (STA or AR), to also evaluate if changes in the TCR immune repertoire persisted or evolved after engraftment and at acute rejection. The patient characteristics are summarized in **Table 1**. The study design is summarized in **Figure 1**. The recipients’ average age was 35±19 yrs. There were 117 males and 83 females. There were 21.5% living related donors, 26% living unrelated donors, and 52.5% deceased donors. All patients received induction with either thymoglobulin (52%) or simulect (48%), and maintenance immunosuppression was low dose maintenance steroids (5mg/day), tacrolimus (target maintenance trough levels after 6 months at 5-8 ng/ml), and mycophenolate mofetil.

**Figure 1.**
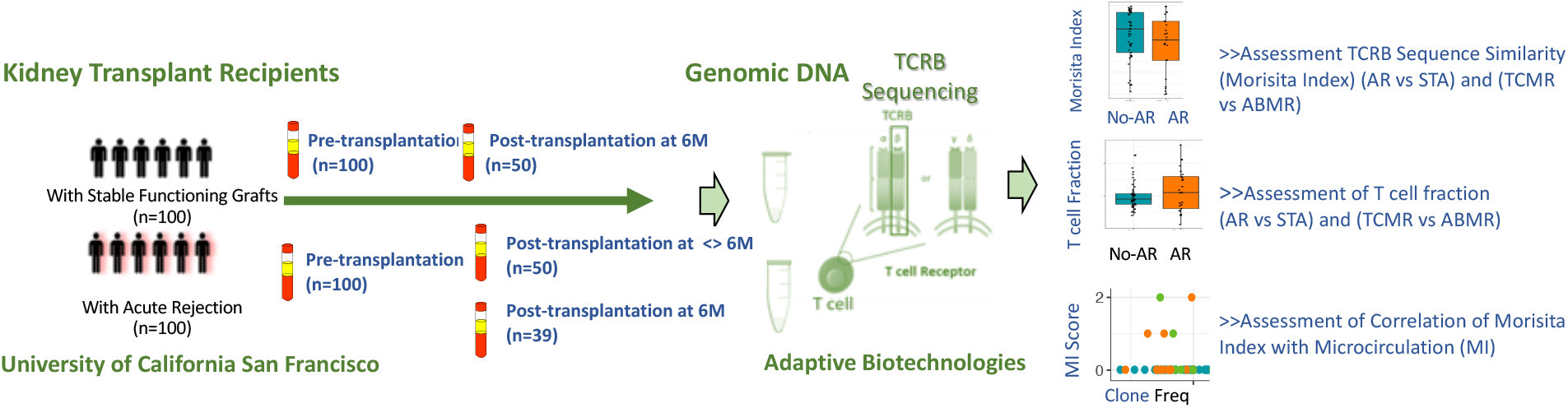
Study Overview. We assessed the immune repertoire of kidney transplant patients and their correlation with AR through TCR sequencing.

### Diversity of the T-cell repertoire and is association with transplant clinical phenotypes

Greater diversity of the T cell repertoire was observed in peripheral blood samples collected from subjects before engraftment, who developed were in the AR patient cohort, and developed acute rejection after transplant. Using a previously published method [21], we inferred the CMV status of the subjects at baseline however it was not able to differentiate between latent versus active infection. A significantly larger number of patients in the AR group were CMV naïve when compared to the STA group (51% vs.26%; Fisher exact P=0.0005). Also, a higher T cell clonality was seen in patients that were inferred CMV PCR positive (n=116) after transplant. (**Figure 2B**). We observe that there is an increase in the CMV positivity rate in the AR group (57% CMV positivity at the time of acute rejection vs 32% CMV positivity in the same patients pre-transplant; p= 0.03). Changes in T-cell repertoire clonality were observed with expansion of specific memory T-cell clones [24], in peripheral blood samples from subjects with biopsy-confirmed acute rejections, with a greater propensity for lower T cell clonality in blood samples collected from paired biopsies with late (> 6 months post-transplant) acute rejection (**Figure 2A**).

**Figure 2.**
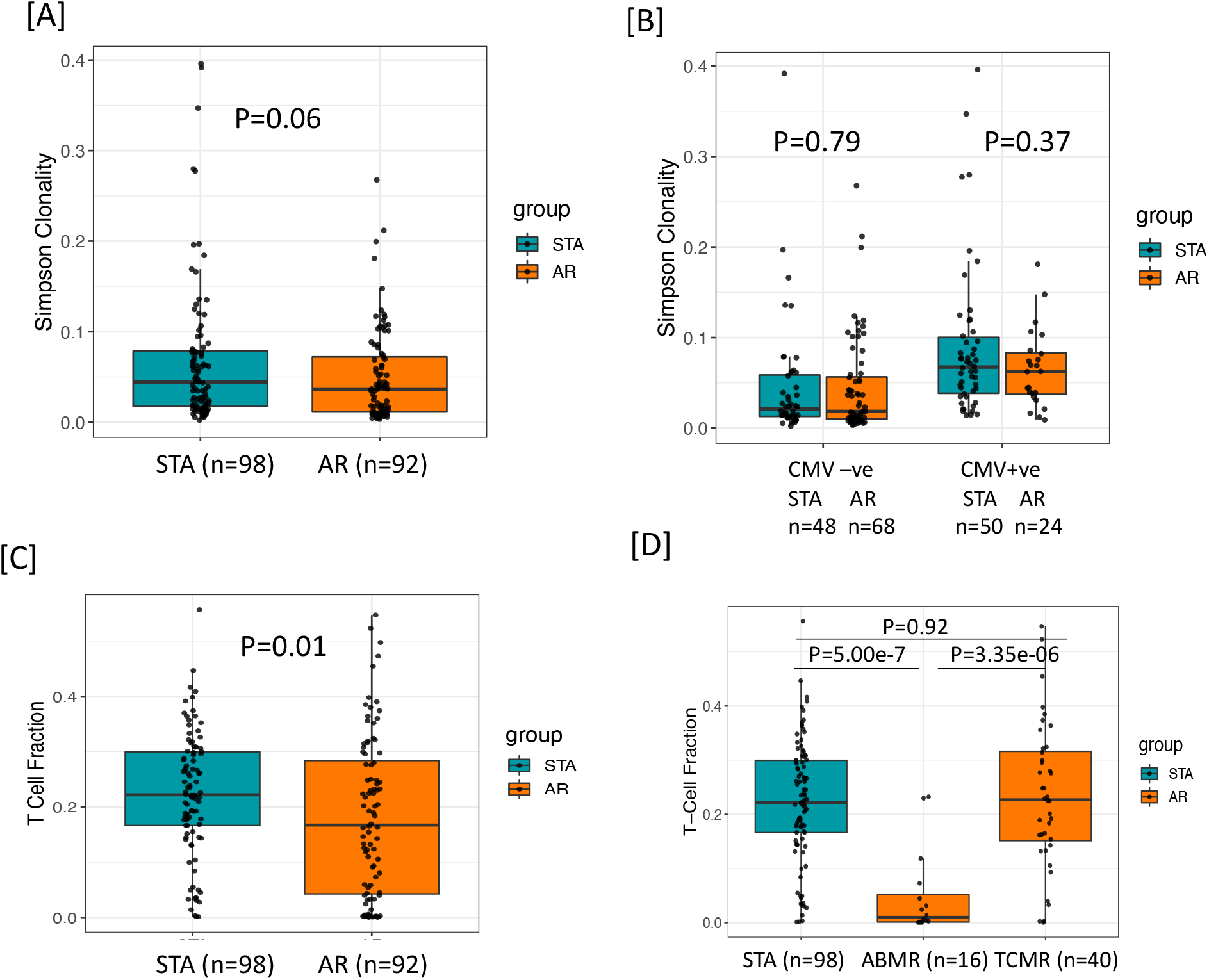
Baseline characteristics of T-cell clonality and TCFr at baseline for transplant recipients who have future rejection and no rejection. [A] There was a trend towards higher clonality in STA subjects, which might indicate a more diverse repertoire at baseline in subjects who will experience AR. [B] There was no difference in baseline repertoire clonality within the groups with or without CMV positivity.[C] At baseline, subjects with low TCFr were at increased risk of future AR (p=0.01). [D] When we analyzed for T-cell fraction among clean TCMR and ABMR cases, we observed a significant difference between the two types of rejection cohorts (Kruskal wallis p=0.026) with a lower value in ABMR relative to TCMR using a post-hoc Dunn test p=0.038.

### Variations in pre-transplant TCRBV01 and TCRBJ01-02 and post-transplant clinical associations

We further assessed the patterns of V and J TCR gene usage within the baseline samples to identify differential gene usage, which may be associated with increased risk of AR (**Supplemental Figure 1A, B**). We observed increased use of TCRBV01 and TCRBJ01-02 in subjects with AR (p<0.05, Wilcoxon rank sum test). Overall, the magnitude of the difference may not be sufficient alone to be predictive of outcomes and may as it may also reflect variations in HLA types among individuals in this data set enriched on patients who reside on the US pacific west coast, this observation requires further validation.

### Baseline TCFr before transplantation and risk of graft rejection

TCFr was determined as the number of T cells out of the total number of nucleated cells in the sample (similar in concept to an estimation of CD3+ T cells by flow sorting)[19]; this denotes the percentage of T cells among all nucleated cells in the blood. We found that subjects with lower TCFr at baseline were at increased risk of rejection and enriched in the AR cohort (p=0.01) (**Figure 2C**). Within the subjects in the AR cohort, no correlation between baseline TCFr and time to rejection was observed (spearman correlation, rho=-0.1 p=0.34). When we analyzed the pretransplant T-cell fraction among patients who had biopsy-confirmed acute rejection classified by Banff [14] as either T cell-mediated rejection (TCMR) or antibody-mediated rejection (ABMR) cases, we observed a significant difference between the two types of rejection cohorts (Kruskal wallis p=4.03e-06) with a lower TCFr value in ABMR using a post-hoc Dunn test p=3.35e-6 (**Figure 2D**). We performed multivariable confounder analysis against the baseline TCFr, and did not observe significant differences for gender, race or recipient age. We observed a sharp rise in TCFr in the AR cohort, specifically for patients who had late acute rejection’s, after the first 6 months post-transplantation (log2-fold change median STA: – 0.34, median <6 months AR: 0.59, median >6 months AR: 3.52; post-hoc Dunn test, adjusted P=0.0037 vs. AR<6mo, P=7.4e-5 vs STA) (**Figure 3A**). When we further stratified rejection patients into ABMR and non-ABMR, irrespective of the time to rejection, TCFr was significantly increased in the ABMR group (AR ABMR vs. STA, Wilcoxon P=0.04, vs. AR non-ABMR vs. STA Wilcoxon P=0.56) (**Figure 3B**). Overall, 70.5% of AR patients had an increase in TCFr post-transplant, compared to 46.9% in STA patients (Fisher exact, p=0.03). In fact, this is because many AR cohort patients had very low baseline TCFr that sharply increased at an average of >10 fold at time of biopsy confirmed rejection. In contrast to TCFr, when we performed a similar analysis for TCR repertoire clonality, we observed no difference between subjects in the AR and STA cohorts (**Supplemental Figure 3),** suggesting that the expansion in TCFr is likely due to an overall increase in the T-cell pool rather than just an expansion of the highest-frequency clones.

**Figure 3.**
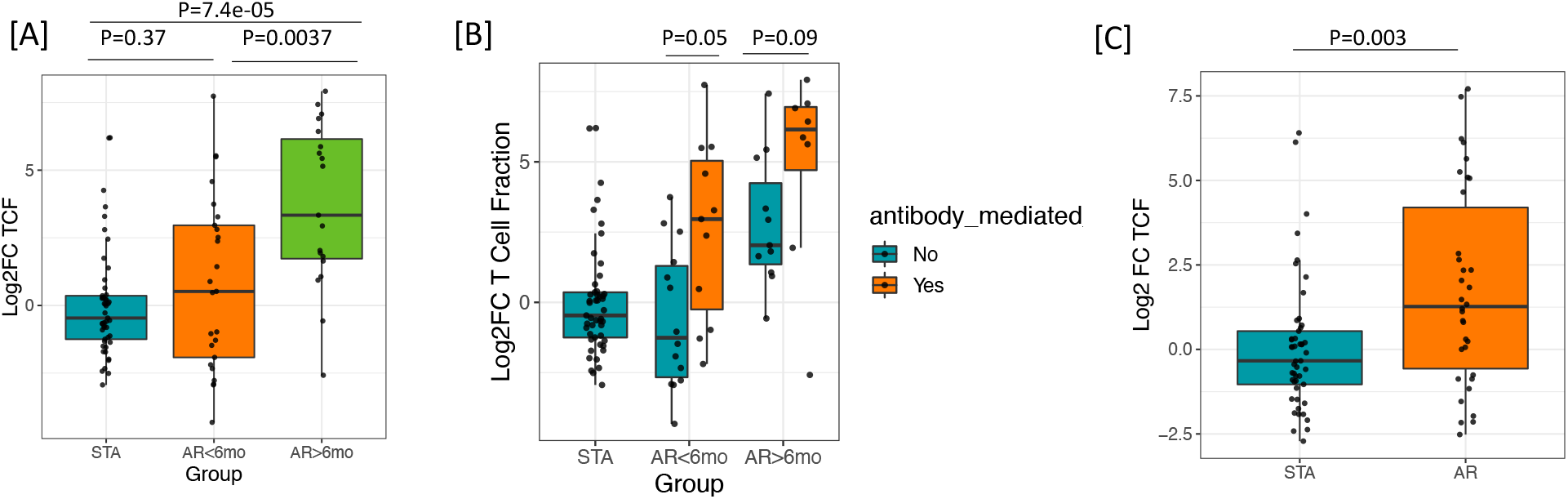
Significant increase in the TCFr at late rejection and in ABMR. (A) An increase of TCFr is significant in late rejection cases >6 months post-transplantation. (B) There is an increased TCFr in ABMR compared to non-ABMR samples. (C) There is an increased peripheral TCFr among AR subjects compared to STA among AR samples collected at 6 months posttransplantation.

### Higher TCR repertoire turnover and acute rejection

T-cell repertoire turnover was measured using Morisita Index [25], which is based on comparing the number and frequency of clones shared between the two samples. We assessed the evolution of TCR repertoire turnover by comparing paired values from the baseline samples and samples collected post-transplant for both STA patients and AR patients, where both samples had greater than 500 T cells. Because repertoire turnover can be influenced by time, we further divided the AR group based on whether their AR event occurred before or after 6 months posttransplant. Increased repertoire turnover was highest in the AR cohort (R=-0.33, P =0.029) (**Supplemental Figure 2**), with highest values for those patients with late rejections after 6 months post -transplant (median Morisita Index for STA: 0.73, median Morisita Index for AR less than 6 months post transplantation: 0.61, median MI for AR post 6 months post transplantation: 0.31, Kruskal-Wallis P=0.004) (**Figure 4A**). To further investigate whether there were clinical characteristics associated with patients that had the highest turnover, we subdivided patients that had ABMR, and found no statistical significance with rejection type (**Figure 4B**). This suggests that high repertoire turnover is generally associated with rejection and not just in patients with antibody-mediated rejection, irrespective of the time to acute rejection (p = 0.0016, adjusted p, post-hoc Dunn test) (**Figure 4C**).

**Figure 4.**
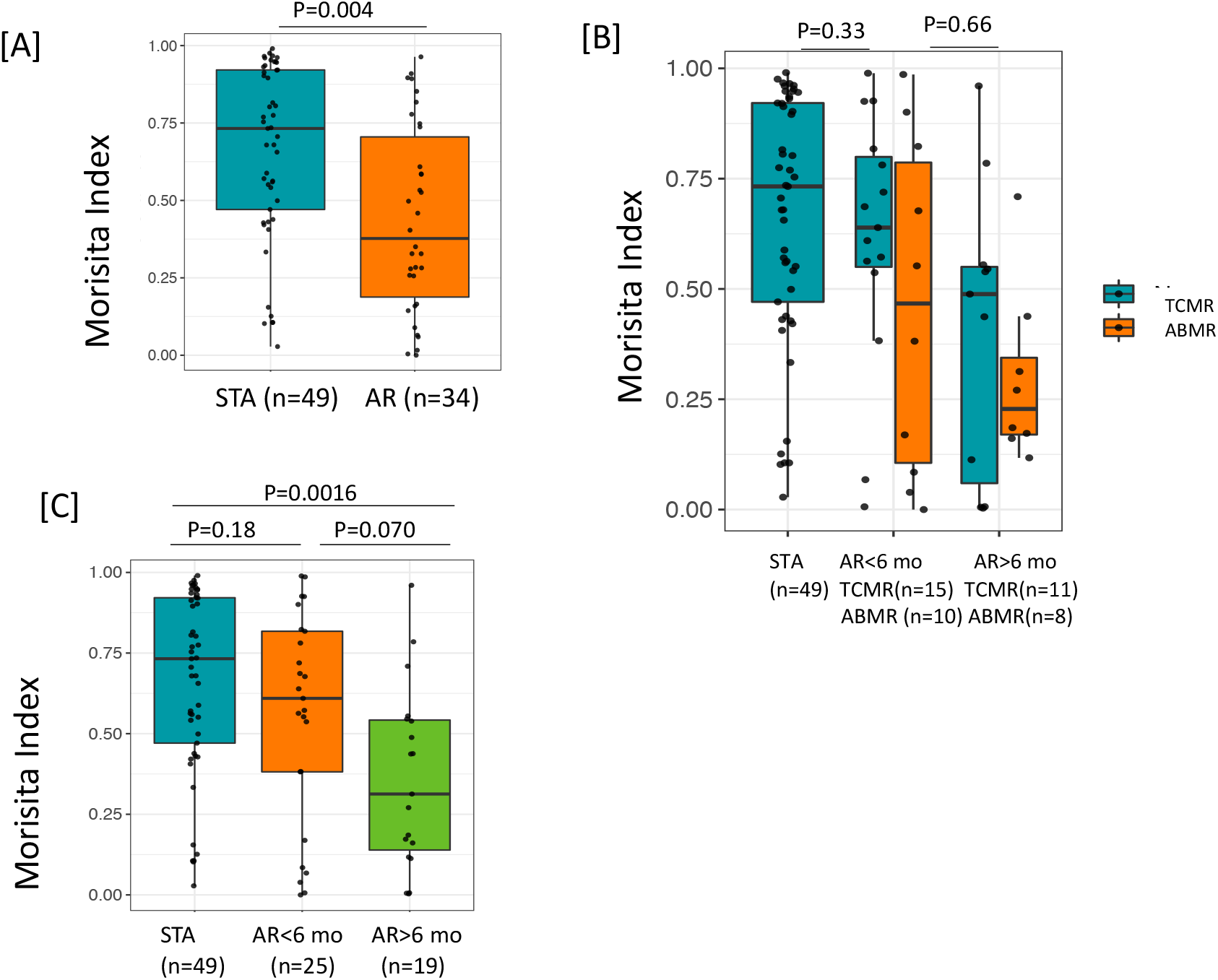
Morisita Index (which measures similarity based on the number of shared sequences between two repertoires and the frequency of overlapping sequences within the two repertoires) was low in AR and in antibody-mediated rejection. (A) The T-cell repertoire turnover is higher (with low Morisita Index) in patients with AR at > 6 mo post-transplantation. (B) T-cell repertoire turnover was more pronounced in ABMR compared with patients with STA and TCMR. (C) There was a greater repertoire turnover at the AR event compared to STA at matching posttransplantation timepoint at 6 mo.

Comparing V and J gene usage post-transplant we observed a slightly greater usage of TCRBV18 in STA patients; however, this was not significant when correcting for multiple testing. No other TCRβ V gene showed differential enrichment (**Supplemental Figure 5A, B**). Together these findings suggest that measuring the TCR immune repertoire changes in an isolated posttransplant sample will likely be insufficient to predict rejection and integrating changes between the pre-transplant or baseline blood sample and a post-transplant blood sample will be more valuable to to evaluate the clinical impact if the repertoire dynamics.

### Microcirculation inflammation score is significantly associated with Morisita Index score

Microcirculation Inflammation (MI), which is a cumulative score that includes glomerulitis (gs) and peritubular capillaritis (ptc), has been associated with worse outcomes in kidney transplantation [26]. A significant correlation was observed between the MI score and Morisita Index in samples taken at post-transplant (spearman’s Rho=-0.27, p=0.0078, **Figure 5**). Of the post-transplant samples with a Morisita index less than 0.3, 43.5% of patients had a MI>0 compared to 15.7% of patients with an MI=0 (Fisher Exact Test, P=0.009). This suggests that greater repertoire turnover might be associated with more severe rejection.

**Figure 5.**
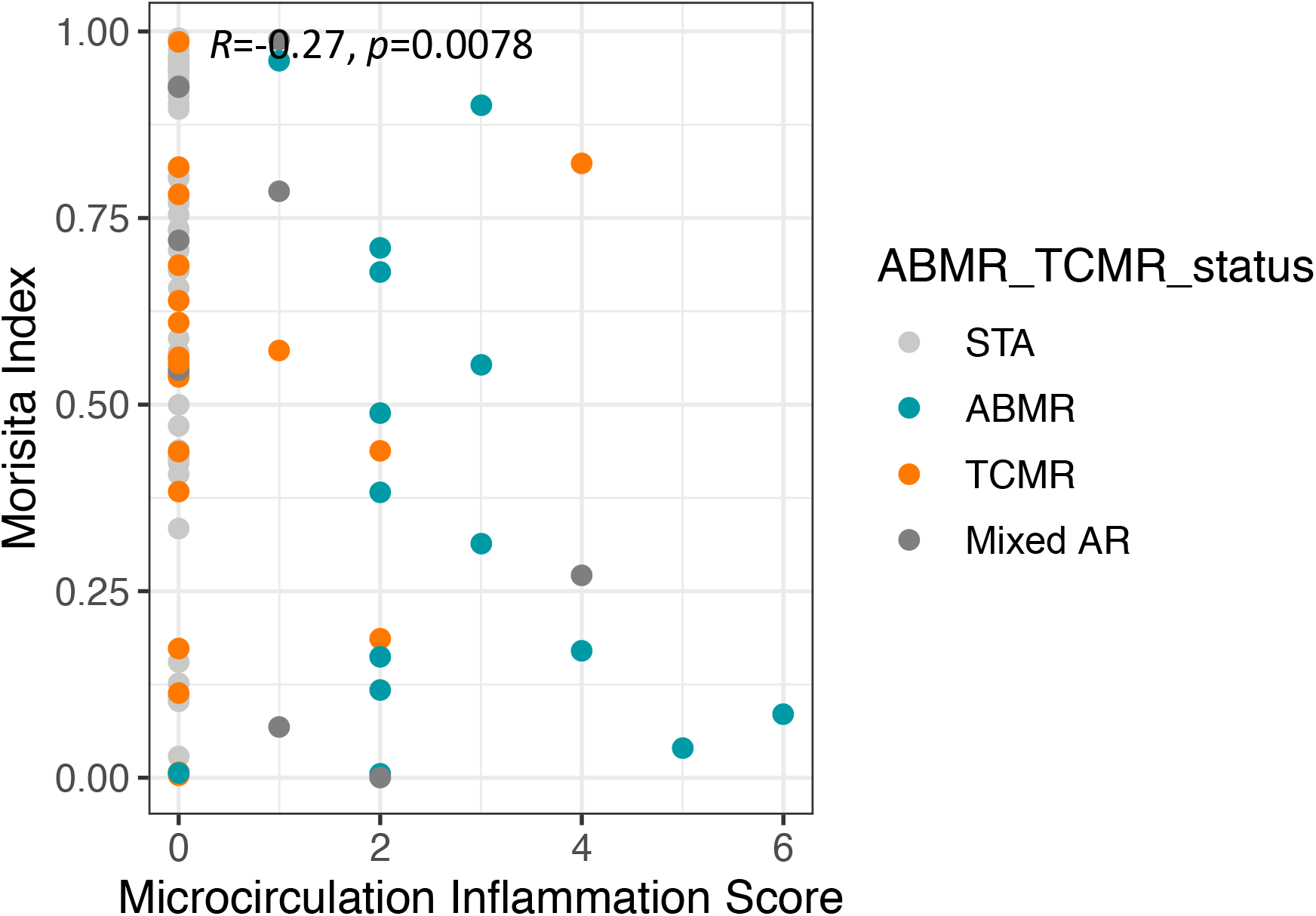
[LEGEND] The Morisita Index inversely proportional to microcirculation inflammation. The finding of increased TCR turnover in terms of low Morisita Index among AR (ABMR, TCMR, Mixed AR) patients was further demonstrated by a significant correlation.

## DISCUSSION

This study provides data on TCR immune repertoire sequencing on the largest cohort of kidney transplant recipients to date, to evaluate the dynamics of TCR changes both before and after transplantation, by analysis of peripheral blood samples paired with transplant biopsies, and paired peripheral blood samples from the same transplant patients, before and after engraftment, using highly validated and demographically matched STA and AR cohorts. Using sequencing of the third complementarity-determining region (CDR3) of the TCRβ chain of the T-cell receptor from 323 blood samples collected from 200 kidney transplant recipients, we report multiple significant observations including associations between AR and Morisita Index and TCFr. Patients with acute rejection had lower (TCFr) before transplantation, followed by greater repertoire turnover and increased TCFr at the time of acute rejection after transplantation. Repertoire clonality is largely driven by the highest frequency clones, thus while there is significant repertoire turnover in patients with rejection, the observed lack of change in clonality indicates the acute rejection is driven by global changes in the TCR repertoire and not specifically by an increase in the highest frequency clones, which are most likely donor reactive. We observe that same repertoire of immunoreactive clones appear to continue to be immunodominant during the post-transplant rejection, even though there is repertoire turnover with regards to clone frequency. Given this observation, one of the reasons for the lower TCFr association with higher risk of post-transplant rejection can be hypothesized to be due to heterologous immunity where cross reactivity with pre-transplant auto-antigens results in increased reactivity to donor allo-specific antigens after transplant, and the evolution of rejection.

Our cohort had a an almost two-fold higher proportion of CMV-recipients pre-transplant in the rejection group (51% vs.26%), as predicted by TCRSeq. We recognize that CMV-recipients overall have a higher risk of CMV infection after transplant, and that there is an increase in the CMV positivity rate in the AR group. As there are known associations between clinical[22] and sub-clinical[23] CMV infection and acute rejection, these observations are relevant. We hypothesize that recipient CMV serostatus may be an independent factor that can influence acute rejection outcomes through sub-clinical CMV viral replication. Nevertheless, though TCRSeq can identify patients with past or current CMV infection, it cannot identify differences between active and latent infection.

TCFr was significantly increased in late rejection cases >6 months post-transplantation and in ABMR. We did not observe significant differences in baseline or post-transplant metrics as single time point metrics, demonstrating the importance of longitudinal tracking. The most significant dynamic trends are observed in ABMR patients with low baseline TCFr that undergo large increases in TCFr and high repertoire turnover post-transplant. This might result from an expansion of CD4^+^ T cells, otherwise observed in ABMR [27–29], but confirmation of this will require further investigation, such as fluorescence-activated cell sorting (FACS) or CyTOF to evaluate the relative contributions of helper and cytotoxic T cells in these rejection episodes with sharp increase in TCFr. The Morisita Index was inversely proportional to MI, and had the ability to predict the mechanism of rejection, i.e. T cell versus antibody-mediated rejection, though this association was expected. These observed associations of TCR immune repertoire changes and late rejection and ABMR, may be explained by the chronicity of allo-immune exposure, the reduction in immunosuppression over time with reduction of immunosuppression target trough levels after 6 months post-transplant, as well as greater intensity of alloimmune injury in ABMR.

Despite the enormous amount of knowledge and data gained form this study, there are some intrinsic limitations as the data was generated on peripheral blood, and we did not have access to profiling paired kidney tissue and/or the draining lymphoid tissue where the immune response could be different or more amplified. Further studies will be required to evaluate change sin the immune repertoire specific to the recovery of T cells following T cell deplting induction such as thymoglobulin, and the inter-individual variability of T cell recovery and TCFr. Further independent validation studies with prospective and serial immune repertoire analysis of the TCR are warranted to validate the clinical utility for of serial TCRSeq for ABMR and late onset acute rejection.

In conclusion, this study validated that detecting repertoire changes correlate with posttransplant rejection episodes suggesting that T-cell receptor sequencing [10] may provide recipient pre-transplant and post-transplant predictors of rejection risk. This assay has the potential to be utilized in immunosuppression management decisions at and after engraftment, based on dramatic increases in TCRf and/or repertoire turnover, where the identification of highcell repertoire turnover post-transplant suggests the expansion of donor antigen-restricted, alloreactive T cells. Additional work is underway to independently validate these results in new patient sets to further decipher T-cell antigen recognition donor-stimulated proliferation of recipient cells to further understand the clonotypic and turnover states that will help to identify dominant epitopes for prediction of renal transplant rejection risk prior to clinical deterioration of graft function.

## Abbreviations

(AR): acute rejection,
(ABMR): antibody-mediated rejection,
(CDR3): complementaritydetermining region,
(CMV): cytomegalovirus,
(MI): microcirculation Inflammation,
(STA): stable,
(TCFr): T-cell Fraction,
(TCMR): T cell mediated rejection,
(TCR): T-cell receptor

## Acknowledgments

We acknowledge the kidney transplant recipients who consented to participate in this study. This study was supported by a National Institutes of Health grant U19 AI128913. We acknowledge Dr. Zoltan Laszik (UCSF) for pathology scores and Dr. Priyanka Rashmi for critically reading the manuscript.

## Author Contributions

MMS, HR, TS were involved in study design, TS, JL, ID, PT, MK, JH, PF, MMS, and MS, contributed to data generation and analysis. MMS, TS, MS, PF, MK contributed to manuscript writing.

---

**Supplemental Figure 1.**
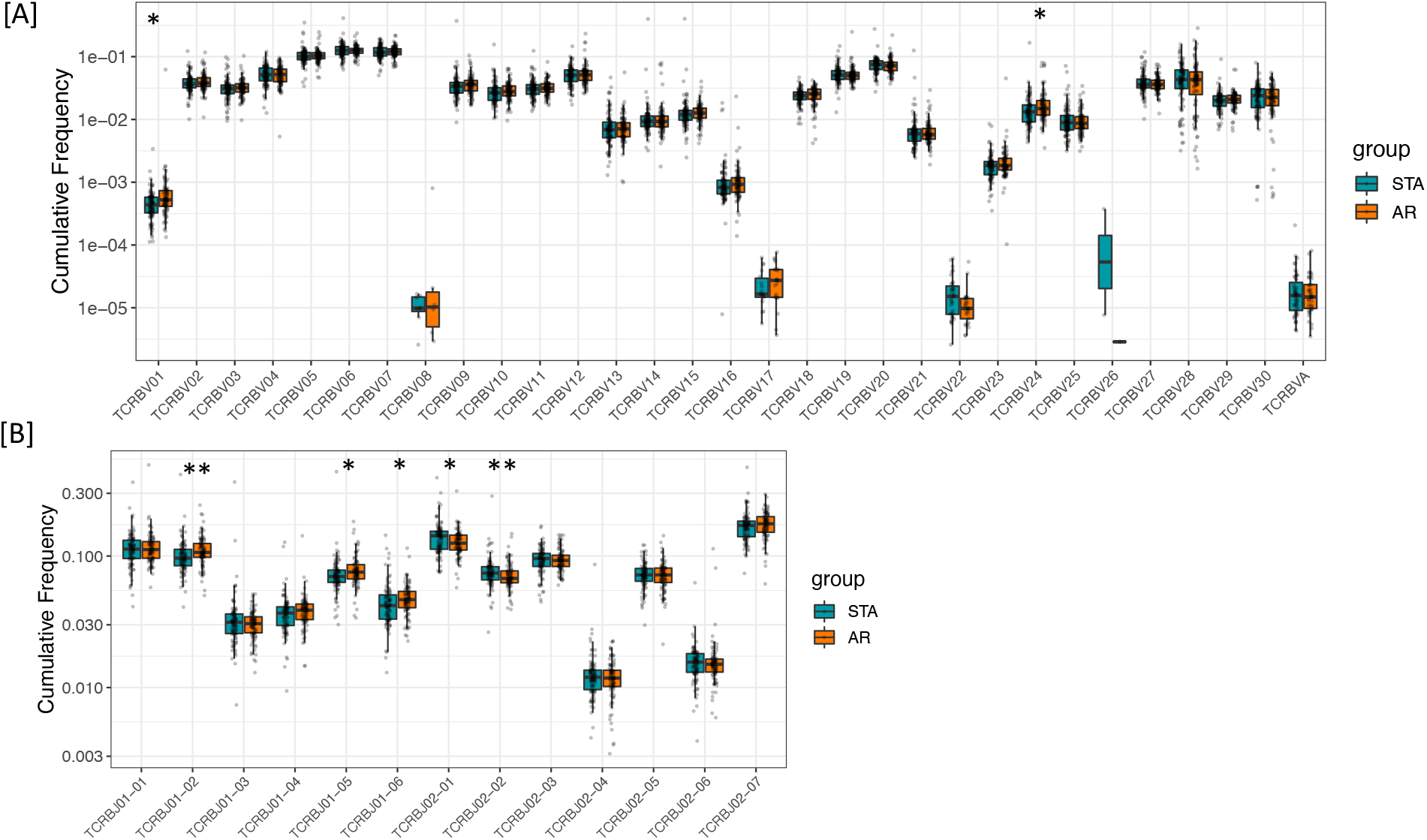
V and J Gene Usage Among Baseline Samples. Cumulative frequency of V [A] and J [B] genes among the 92 AR and 98 STA baseline samples. P values determined by Wilcox rank sum test, * p<0.05, ** p<0.01.

**Supplemental Figure 2.**
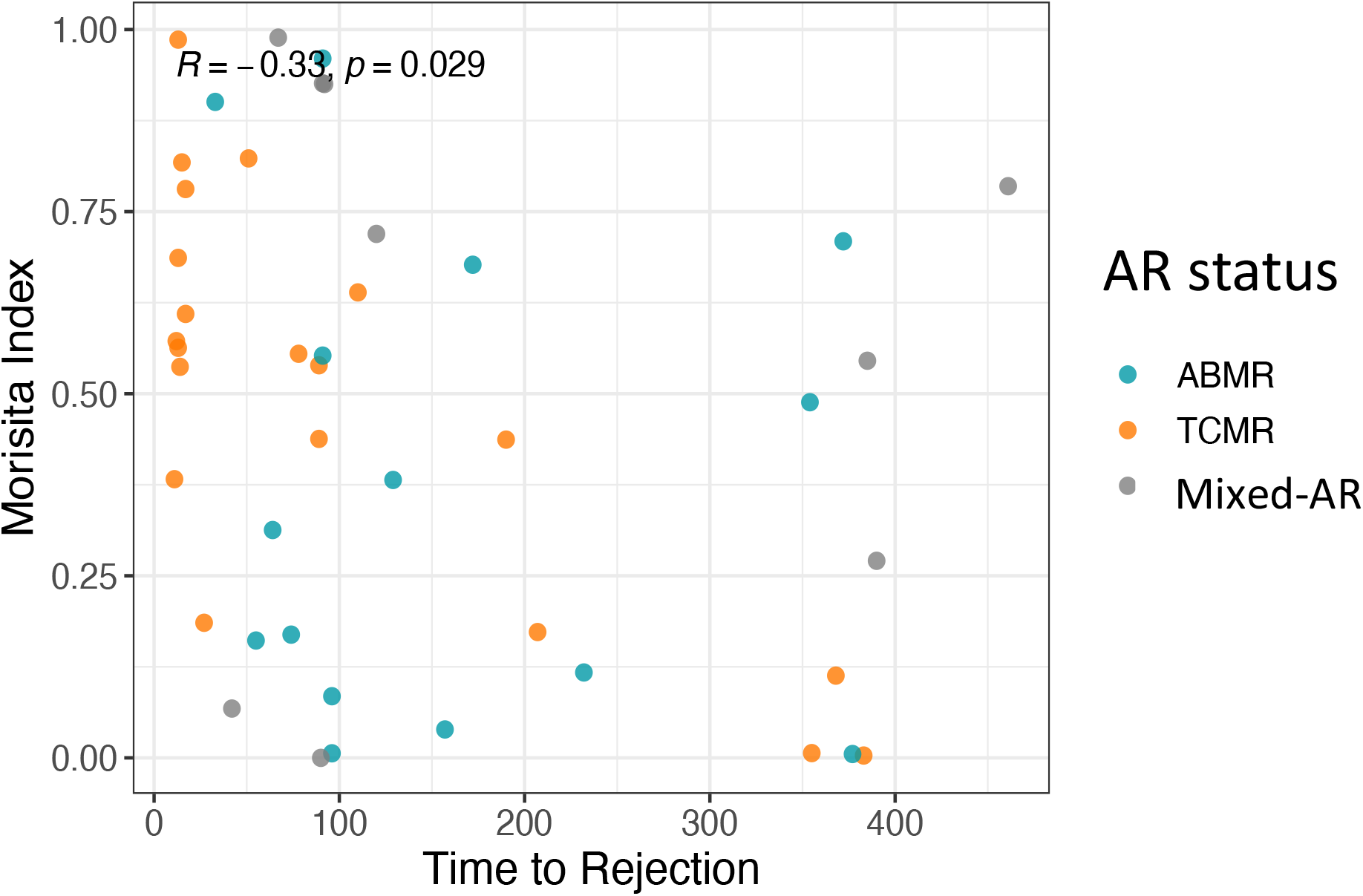
Morisita Index correlates with post-transplant time until AR episode. Among the AR cohort (ABMR, TCMR, and mixed AR included that had features of both TCMR and ABMR), Morisita Index was significantly correlated with time to AR with spearman’s Rho, R=0.33, P=0.029.

**Supplemental Figure 3.**
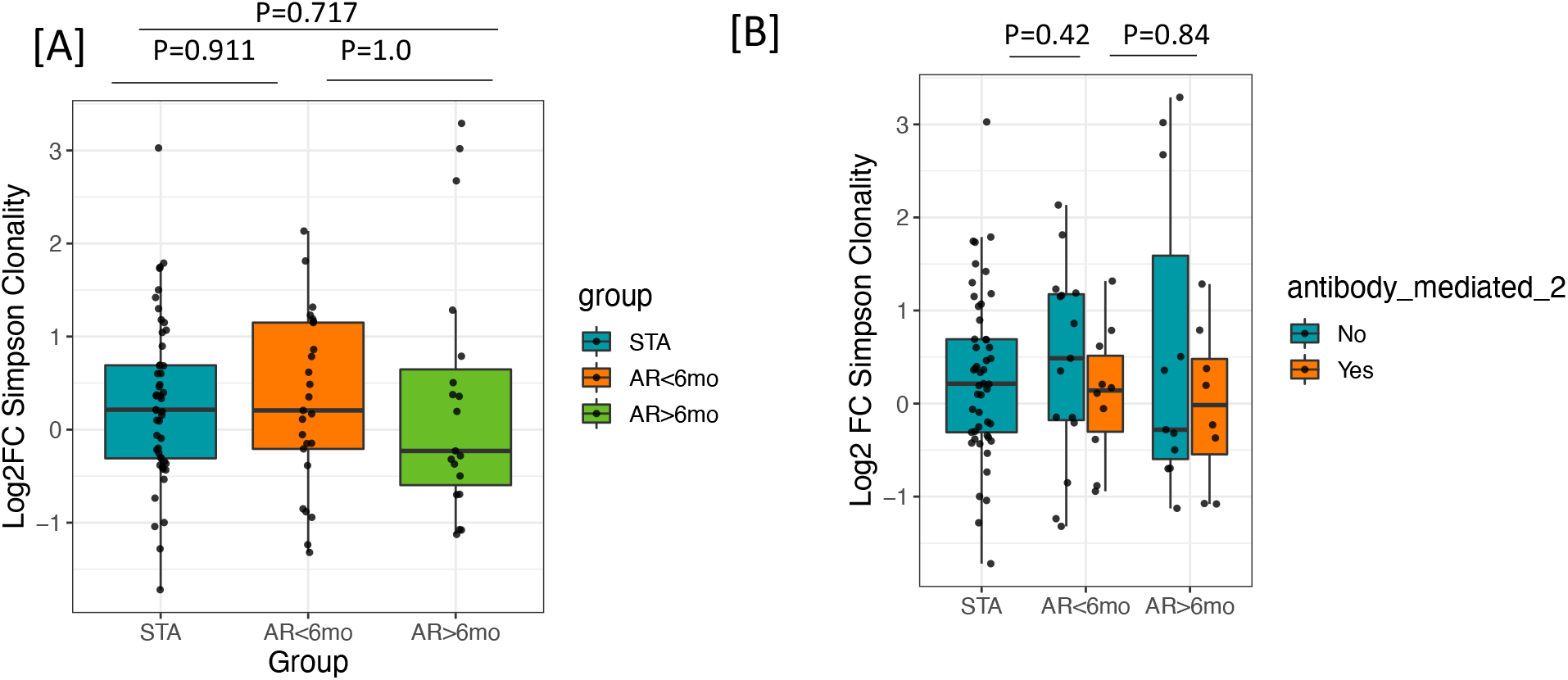
Change in Repertoire Clonality Is Not Associated with AR. (A) Patients with AR do not show a significantly different change in repertoire clonality relative to STA patients. (B) There is no significant difference in repertoire clonality in ABMR compared to non-ABMR samples.

**Supplemental Figure 4.**
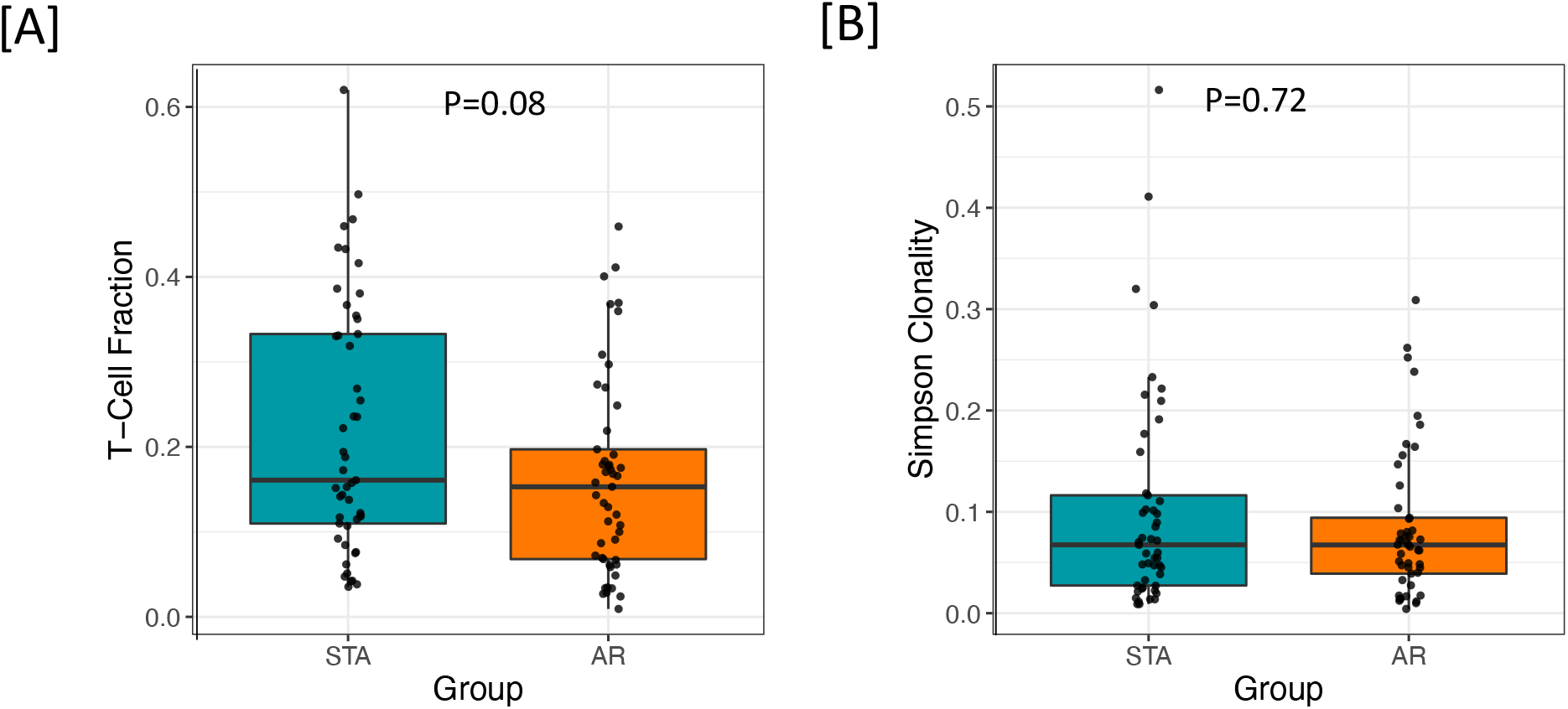
Post-Transplant Repertoire Does Not Vary Between STA and AR Patients.

**Supplemental Figure 5.**
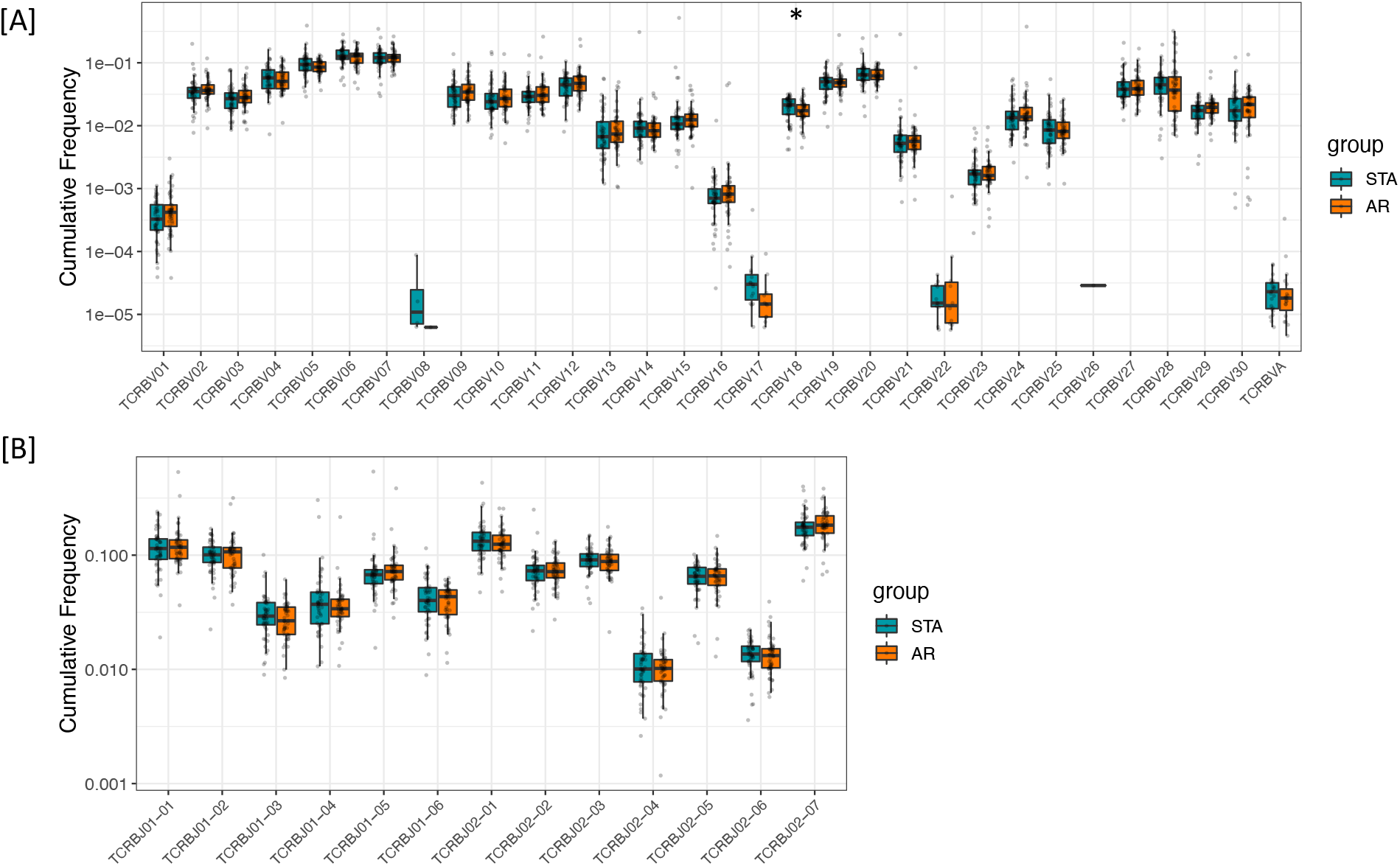
V and J Gene Usage Does Not Vary Post Transplant. Cumulative frequency of V [A] and J [B] genes among the 49 AR and 49 STA Post-transplant samples. P values determined by wilcox rank sum test, * p<0.05.

